## Supplementary Information for "Single-molecule digital sizing of proteins in solution"

**Analysis of diffusion profiles using the advection–diffusion model**

**Overview.** To extract the hydrodynamic radius $R_H$, we analyze the experimental diffusion profiles using an advection–diffusion model. This involves fitting the experimental profiles with a sequence of simulated profiles, referred to as basis functions, generated through numerical simulations. By applying a least-squares error algorithm, we identify the simulated profiles that best corresponds to the experimental data, which enables us to extract the diffusion coefficient $D$. Subsequently, $R_H$ is deduced using the Stokes–Einstein equation.

**Generation of basis functions.** The basis functions are generated following a method described by Q.A.E. Peter.[1] These functions are derived from a dimensionless differential equation rooted in the advection–diffusion equation, which describes the diffusion of species in a Poiseuille flow within a rectangular microfluidic channel. The model considers the diffusion of particles in a steady-state flow with a constant flow rate $Q$ within a channel of height $H$ and width $W$.

Basis functions are derived by first considering the general advection–diffusion equation for the local concentration $c(x, z, y)$ of the species within the channel. This equation accounts for the combined effects of advection due to fluid flow and molecular diffusion. The advection–diffusion equation is given by:

$$\frac{\partial c}{\partial t} = \nabla \cdot (D\nabla c) - \nabla \cdot (\vec{v}c) \tag{S1}$$

with $D$ being the diffusion coefficient, $\vec{v}$ the flow velocity, and $t$ the time.

Under the assumptions of steady-state conditions (i.e., time invariance) and the presumption that diffusion along the channel is negligible (i.e., flow in $x$-direction is much stronger than diffusion), Equation S1 simplifies to:

$$\partial_x c = \frac{D}{v_x}\left(\partial_y^2 + \partial_z^2\right) c \tag{S2}$$

This expression can be reformulated into a dimensionless form:

$$v_x' \partial_\Phi c = \left(\partial_{y'}^2 + \partial_{z'}^2\right) c \tag{S3}$$

with $\Phi$ being the dimensionless "diffusion coefficient", expressed as $\Phi \equiv x\frac{D\beta}{Q}$, where $x$ is the distance from the inlet, and $\beta$ denotes the aspect ratio, which is given by $\beta = \frac{H}{W}$. The variables $y'$ and $z'$ are the dimensionless axes coordinates, with $y' \equiv \frac{y}{W}$ and $z' \equiv \frac{z}{H}$, respectively. The dimensionless velocity $v_x{}'$ (for specific $y', z'$ positions) is represented as $v_x{}' \equiv v_x\frac{WH}{Q}$.

$v_x{}'$ can be determined from the incompressible Navier–Stokes equation. This equation links the fluid's pressure $p$, viscosity $\eta$, and density $\rho$ to its velocity vector $\vec{v}$ within the channel, and provides a way to compute the velocity profile in the microchannel:

$$\rho\left(\partial_t\vec{v} + \left(\vec{v}\cdot\vec{\nabla}\right)\vec{v}\right) = -\vec{\nabla}p + \eta\nabla^2\vec{v} \tag{S4}$$

By applying the principles of time invariance and translational invariance along with the dimensionless parameters defined earlier, Equation S4 can be reformulated as:

$$\left(\partial_{y'}^2 + \partial_{z'}^2\right) v_x'(y', z') = 1 \quad \text{(S5)}$$

Equation S5 can then be solved using a series expansion method. Specifically, the velocity $v_x'(y', z', \beta)$ is given by the series:

$$v_x'(y', z', \beta) = \sum_{nm} A_{nm}(\beta) \sin(n\pi y') \sin(m\pi \frac{z'}{\beta}) \quad \text{(S6)}$$

where the coefficients $A_{nm}(\beta)$ are defined as:

$$A_{nm}(\beta) \equiv -\frac{16}{\pi^4} \frac{1}{nm\left(n^2 + \frac{m^2}{\beta^2}\right)} \delta_{(n \bmod 2),1} \delta_{(m \bmod 2),1} \quad \text{(S7)}$$

Here, $\delta$ denotes the Kronecker delta function and the variables $n$ and $m$ are integers used in the summation, representing the series terms for the horizontal $y'$ and vertical $z'$ dimensions, respectively. This formulation allows to calculate the velocity profile within the channel.

Using $v_x'$ from Equation S6, we can now numerically integrate Equation S3 using the Crank–Nicholson method. This approach enables us to construct diffusion profiles at specific distances $x$ within the channel, which correspond to various measurement positions, for a range of diffusion coefficients $\Phi$ (representing different $R_H$ values). These generated profiles serve as the basis functions $B(\Phi)$.

At each measurement position, a set of these basis functions $B(\Phi)$ is generated, with each representing a distinct value of $\Phi$ (or different diffusion coefficients $D$). We typically use a set of 1000 logarithmically spaced $\Phi$ values, encompassing hydrodynamic radii from 0.1 nm to 1 µm. The initial concentration profile, used to compute the initial conditions for the simulations, is derived from the first fluorescence profile measured. We assume that this concentration is uniformly distributed throughout the height of the channel ($c_i(y, z) = c_i(y)$).

To ensure accurate and consistent representation across various diffusion coefficients, we normalize the basis functions $B(\Phi)$. This normalization accounts for potential variations in signal amplitude and baseline offset that may occur due to differential illumination across different images, ensuring consistent and accurate representation across the range of diffusion coefficients.

**Fitting procedure.** The process of fitting involves calculating the least square error between the experimental diffusion profiles and the simulated basis functions $B(\Phi)$ for each value of $\Phi$. The fit is conducted globally across all measurement positions in the microchannel. To determine the optimal fit, we identify the two best matching basis functions $B(\Phi)$ using the sum of squared errors (SSE). These are then interpolated to ascertain the fitted diffusion coefficient $\Phi$. The hydrodynamic radius $R_H$ is subsequently calculated using the Stokes–Einstein relation, given by the formula:

$$R_{\mathrm{H}} = \frac{x}{Q\Phi} \beta \frac{k_B T}{6\pi\eta} \quad \text{(S8)}$$

where $x$ is the distance from the inlet, $Q$ is the flow rate, $\Phi$ is the dimensionless diffusion coefficient, $\beta$ is the aspect ratio of the channel height to its width, $k_B$ is the Boltzmann constant, $T$ is the absolute temperature, and $\eta$ is the fluid's viscosity.

**Error calculation.** The variance $\sigma_\Phi^2$ is calculated using the first-order Taylor series of the least squares equation, which is given by:

$$\sigma_\Phi^2 = \frac{1}{\sum_i \left(\frac{\partial B_i(\Phi)}{\partial \Phi}\right)^2 \frac{1}{\sigma_i(\Phi)}} \tag{S9}$$

with $\sigma_i$ being the noise on each point *i* of the profile. Due to the non-symmetric shape of the SSE function around the determined $\Phi$, it is not appropriate to report errors as simple symmetric $\pm\sigma_\Phi^2$ values. To address this asymmetry, we first identify the side exhibiting the greater SSE, either $SSE(\Phi - \sigma_\Phi^2)$ or $SSE(\Phi + \sigma_\Phi^2)$, which subsequently determines the noise level. We then calculate the $\Phi$ values at which this noise level intersects with the SSE function, thus defining the error range for $\Phi$. The two limits of the range $\Phi$ is then converted using Equation S8 to get the error range for $R_H$.

**Analysis code.** Analysis of diffusion profiles using the advection–diffusion model is done via a custom-written code in Python. The computational analysis code, developed by Q.A.E. Peter, is available on the Zenodo repository: https://doi.org/10.5281/zenodo.3881940. The current version is available on https://github.com/impact27/diffusion_device/tree/v1.0.0. For our study's specific needs, we created a dedicated branch of this original code, which includes tailored examples and instructions for analyzing traces from smMDS experiments. This branch is accessible via the GitHub repository: https://github.com/gkrainer/smMDS.

### Supplementary Figures

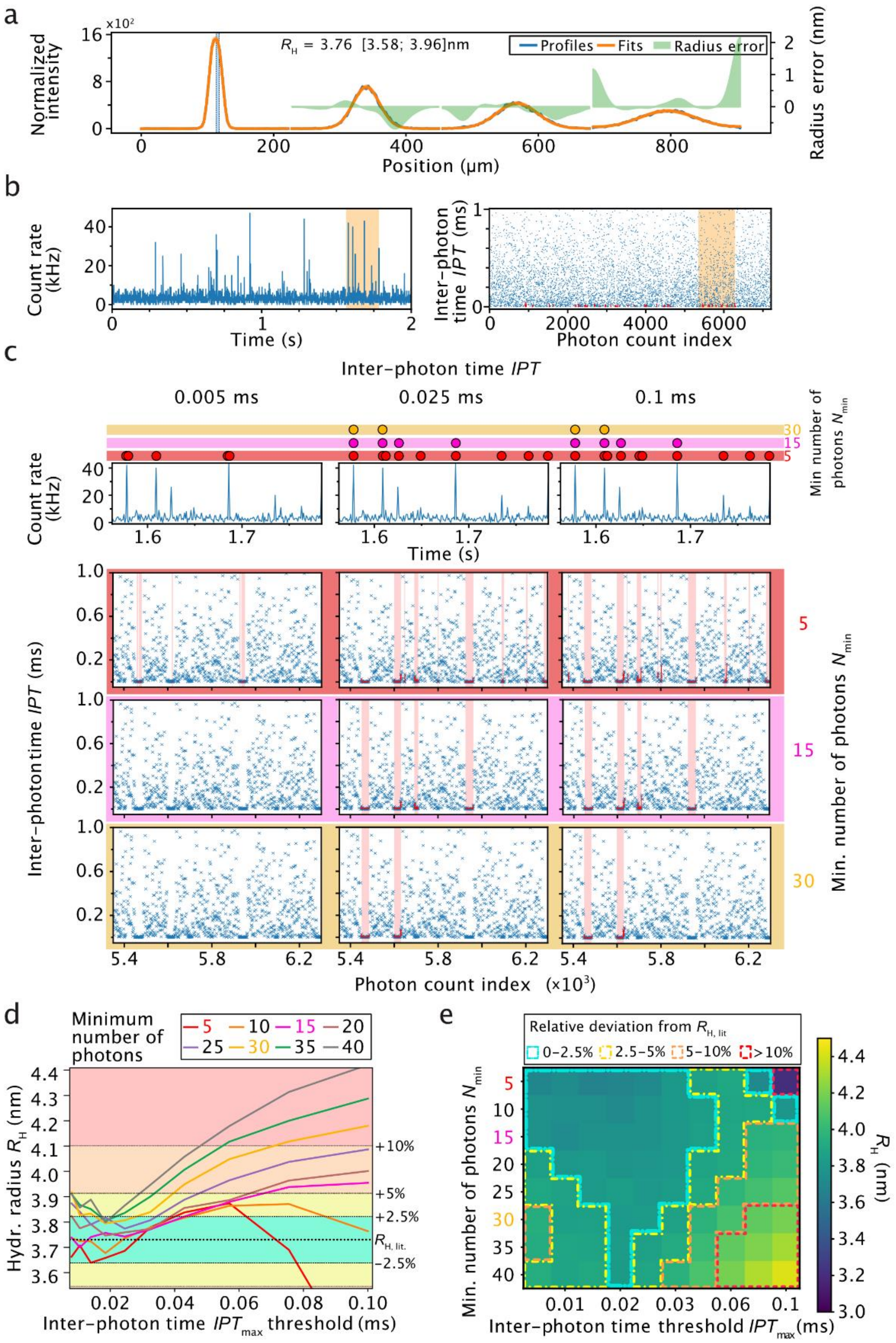


**Supplementary Figure 1. Influence of burst parameter selection in smMDS experiments. (a)** Diffusion profile for HSA as obtained from a step scan measurement at 20 pM protein concentration. Diffusion profile was generated from 2-second-long step scans across four channels of the microfluidic chip. Burst selection parameters used to obtain the diffusion profile were as follows: maximum inter-photon time $IPT_{max}$ = 0.02 ms, minimum number of photons $N_{min}$ = 15, and Lee filter 4. Diffusion profile is shown in blue, experimental fit in orange, and errors are shown as green bands. The extracted hydrodynamic radius $R_H$ [with errors] is given as an inset. **(b)** Photon-count time trace (left, 1-ms binning) and inter-photon time plot (right) at diffusion profile position 112.52 µm, as indicated with a blue line in panel a. Yellow highlighted regions in the photon-count time trace and the inter-photon time plot highlight the zoom-ins shown in panel c. Red highlighted regions in the inter-photon time plot are photon events within single-molecule bursts. **(c)** Influence of burst parameter selection on detection of single-molecule events. Shown are photon-count time traces (top panel, 1-ms binning) and inter-photon time plots (bottom panel) for different combinations of $IPT_{max}$ ($IPT_{max}$ = 0.005 ms, 0.025 ms, and 0.1 ms) and $N_{min}$ thresholds ($N_{min}$ = 5, 15, and 30) for the yellow highlighted region in panel b. Lee filter was 4 in all cases. Colored dots indicate detected single-molecule events for different $N_{min}$ threshold values (red: $N_{min}$ = 5, pink: $N_{min}$ = 15, and yellow: $N_{min}$ = 30). Red highlighted regions in the inter-photon time plots are photon events within identified single-molecule bursts. **(d)** Influence of burst parameter selection on the size determination of HSA. $R_H$ values were extracted from diffusion profiles generated from different combinations of $IPT_{max}$ and $N_{min}$ values by varying $IPT_{max}$ from 0.01–0.1 ms and $N_{min}$ from 5–40. The dashed line indicates the average literature value for HSA ($R_{H,lit.}$ = 3.73 ± 0.40 nm)[2–5] with cyan, yellow, orange, and red bands denoting 0–2.5%, 2.5–5%, 5–10%, and >10% relative deviations from $R_{H,lit}$ values, respectively. **(e)** Two-dimensional representation of the influence of different $IPT_{max}$ and $N_{min}$ thresholds values on the size determination of HSA. Extracted $R_H$ values for different combinations of $IPT_{max}$ and $N_{min}$ values are color coded (from $R_H$ = 3.0 nm, blue to $R_H$ = 4.5 nm, yellow), see color bar on the right). Regions with 0–2.5%, 2.5–5%, 5–10%, and >10% relative deviations from $R_{H,lit.}$ values are highlighted in cyan, yellow, orange, and red, respectively.

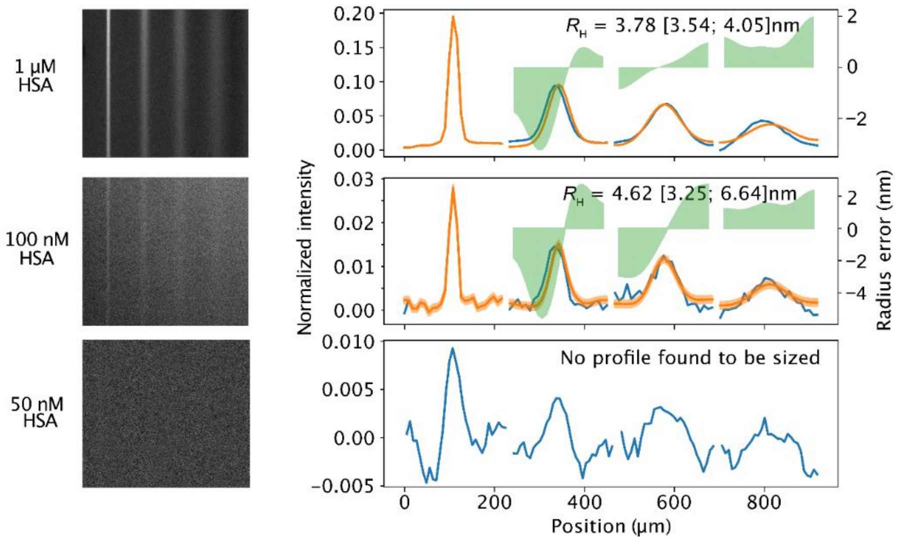


**Supplementary Figure 2. Diffusional sizing of HSA by conventional widefield imaging MDS.** Experimental images from widefield microscopy (left panel) and diffusion profiles (right panel) of HSA taken at different protein concentrations (1 µM, 100 nM, and 50 nM HSA, from top to bottom). Experiments were performed on a custom-built epifluorescence microscope for MDS experiments as described in Schneider et al.[6] The chip was the same as used in smMDS experiments. Diffusion profiles were recorded at four different positions along the microfluidic channel. The flow rates were 100 µL/h and the buffer conditions the same as used in smMDS experiments (PBS supplemented with 0.01% Tween 20). Diffusion profiles are shown as blue lines, experimental fits as orange lines, and errors as green bands. Extracted hydrodynamic radii $R_H$ [with errors] are given as insets. No fit could be obtained for the measurement at 50 nM HSA.

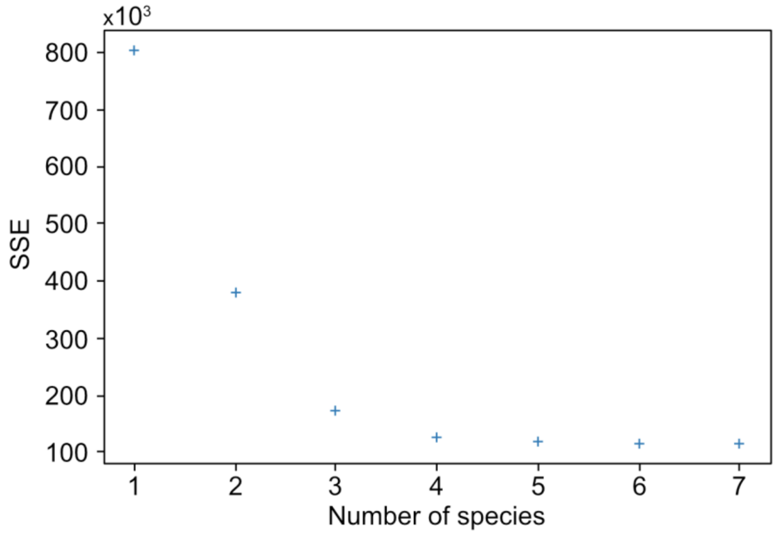


**Supplementary Figure 3: Goodness of fit for different fitting models applied to the burst intensity histogram shown in Figure 5a.** Shown is the sum of square errors (SSE) for each model, which includes one skew distribution and (n–1) Gaussian distributions, where *n* denotes the total number of distributions (corresponding to the total number of species analyzed). The graph demonstrates that beyond four species (monomer to tetramer), there is no visually discernible improvement in the fit.

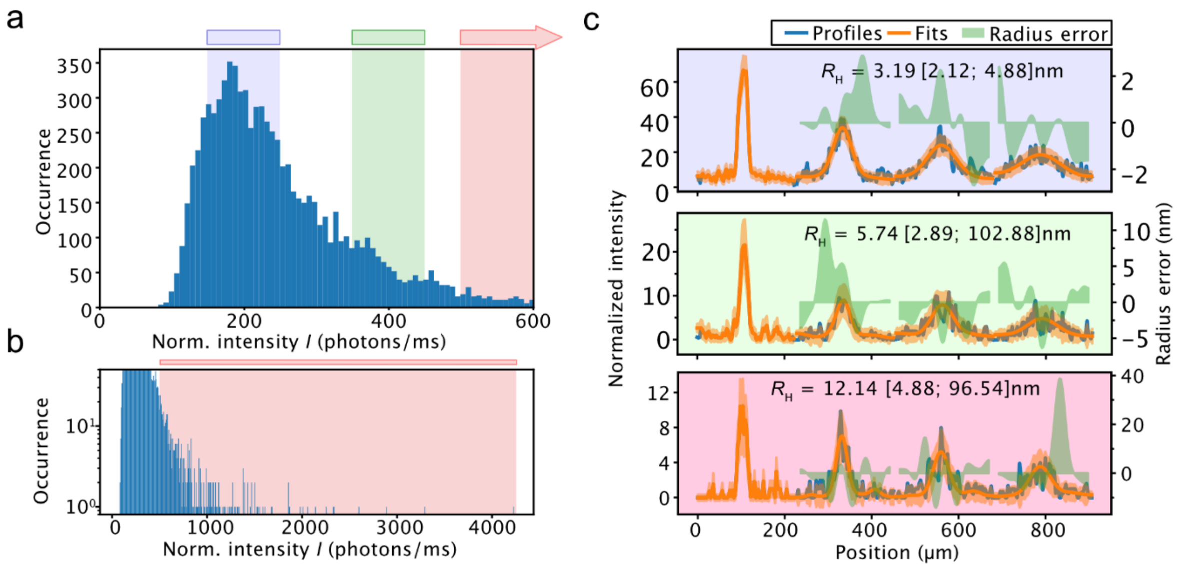


**Supplementary Figure 4. Resolving α-synuclein oligomer populations by selecting defined regions in the burst intensity histogram. (a)** Distribution of α-synuclein oligomer burst intensities. All bursts extracted from a smMDS step scan measurement of α-synuclein oligomers (10 pM) are displayed in a burst intensity histogram. Intensities are normalized intensities with respect to burst duration. Three regions (blue, green, and red) are selected in the burst intensity histogram. The blue and green regions span an intensity region of 100 photons/ms, starting from 150 and 350 photons/ms, respectively. The red region spans from 500 to 4230 photons/ms (indicated by the arrow). The entire red region is shown in panel b. **(b)** Burst intensity histogram in semi-log scale with red region highlighted. **(c)** Diffusion profiles generated from bursts within each of the three regions defined in panel a. The obtained diffusion profiles were fitted to extract size information. Diffusion profiles are shown as blue lines, experimental fits as orange lines, and error as green bands. Extracted $R_H$ [with errors] are given as insets.

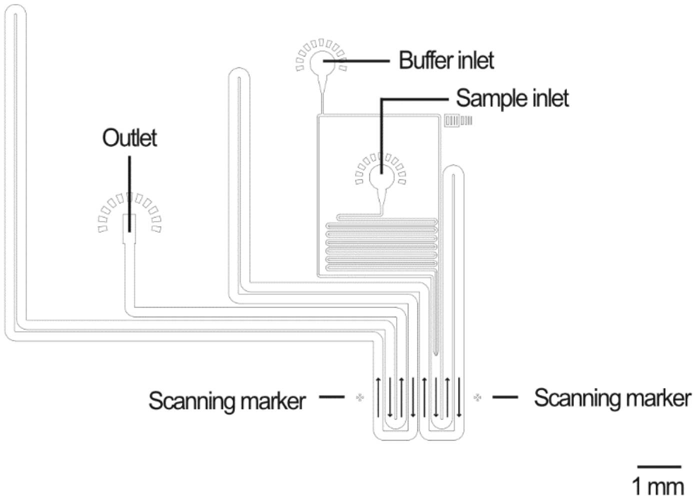


**Supplementary Figure 5. Schematic of the microfluidic chip design.** The most relevant features are highlighted. The arrows indicate the flow path within the channels.

### Supplementary Tables

**Supplementary Table 1. Extracted hydrodynamic radii of HSA as displayed in Figure 2d.**

| HSA concentration | Hydrodynamic radius $R_H$ (nm)[a] Continuous scan | Step scan |
|---|---|---|
| 100 fM | | 3.99 ± 0.31 |
| 1 pM | | 3.73 ± 0.050 |
| 10 pM | | 3.75 ± 0.21 |
| 20 pM | 3.37 ± 0.19 | 3.72 ± 0.035 |
| 50 pM | 3.58 ± 0.11 | |
| 100 pM | 3.66 ± 0.036 | |
| 1 nM | 3.71 ± 0.051 | |
| 10 nM | 3.58 ± 0.037 | |
| 100 nM | 3.39 ± 0.11 | |
| 1 µM | 3.83 ± 0.12 | |

[a] Errors are reported as standard deviations.

**Supplementary Table 2. Extracted hydrodynamic radii of proteins as displayed in Figure 3a.**

| Protein | Hydrodynamic radius $R_H$ (nm)[a] |
|---|---|
| Alexa 488 | 0.60 ± 0.089 |
| Lysozyme | 1.66 ± 0.21 |
| RNase A | 2.37 ± 1.37 |
| α-Synuclein | 3.14 ± 0.37 |
| HLA | 2.86 ± 0.28 |
| HSA | 3.75 ± 0.21 |
| Thyroglobulin | 8.21 ± 0.42 |
| α-Synuclein oligomers | 7.50 ± 1.27 |

[a] Errors are reported as standard deviations.

**Supplementary Table 3. Quantification of protein labeling.**

| Protein | Concentration used for labeling | Molar excess dye | Degree of labeling (DOL) | Concentration after labeling and purification |
|---|---|---|---|---|
| RNase A | 20 µM | 3x | 0.90 | 11.8 µM |
| HSA | 20 µM | 3x | 0.99 | 11.9 µM |
| Lysozyme | 40 µM | 3x | 0.34 | 24.5 µM |
| HLA | 3.12 µM | 6x | 0.24 | 734 nM |
| Thyroglobulin | 6.45 µM | 5x | 0.34 | 2.48 µM |
| α-Synuclein | 120 µM | 3x | 0.67 | 45.6 µM |
